# Circuit-specific sonogenetic stimulation of the deep brain elicits distinct signaling and behaviors in freely moving mice

**DOI:** 10.1101/2021.11.06.467579

**Authors:** Quanxiang Xian, Zhihai Qiu, Shashwati Kala, Kin Fung Wong, Suresh Murugappan, Yong Wu, Xuandi Hou, Jiejun Zhu, Jinghui Guo, Lei Sun

## Abstract

Sonogenetics uses heterologously-expressed proteins to sensitize neurons to ultrasound, enabling selective, non-invasive, and deep brain stimulation. However, its ability to modulate specific circuits or induce behavioral changes remains to be studied and characterized. Here, we demonstrate that sonogenetics enables efficient activation of well-defined neural circuits by transcranial low-intensity, low-frequency ultrasonic stimulation with high spatiotemporal resolution. Targeted neurons in subcortical regions were made to express a mechanosensitive ion channel (MscL-G22S). Ultrasound could trigger activity in MscL-expressing neurons in the dorsal striatum without increased activation in neighboring regions, and increase locomotion in freely-moving mice. Ultrasound stimulation of MscL-expressing neurons in the ventral tegmental area could activate the mesolimbic pathway to trigger dopamine release in the nucleus accumbens and modulate appetitive conditioning. In MscL-expressing cells, neuronal responses to ultrasound pulses were rapid, reversible and repeatable. Altogether, we show that sonogenetics can selectively manipulate targeted cells to activate defined neural pathways and affect behaviors.

## Introduction

It has been a long-standing goal of neuroscience to specifically manipulate neural activity to understand the brain’s functions and dysfunctions. Sonogenetics, analogous to optogenetics, combines the targeted expression of mechanosensitive cellular machinery (e.g. ion channels) and precise delivery of ultrasound (US). This approach has been shown to enable non-invasive stimulation, high spatiotemporal resolution, accurate deep-brain targeting through cell-type-specific gene expression [1–6], and could potentially be translated to application in large mammals, and possibly even humans. ‘Sonogenetics’ was first described in a study showing that ultrasound, in the presence of microbubbles, could activate neurons overexpressing the mechanosensitive ion channel TRP-4 to control the behaviors of *C. elegans* worms [2]. Later studies have used different candidates to successfully enhance neuronal response to ultrasound stimulation, including the mechanosensitive proteins Prestin [3], MscL [1, 7], and hsTRPA1 [6]. The MscL channel in particular is an intriguing candidate as it is a mechanically-activated ion channel that can quickly convert mechanical stimuli into neuronal responses [8], independent of membrane potential [7] and other endogenous factors and thermal deposition [9]. Indeed, our previous work has shown that the expression of MscL can sensitize neurons to ultrasound and specifically induce neuronal activation in the region expressing it [1].

However, a spatiotemporally-coordinated activation of specific neural networks is required to control outcomes beyond the cellular level, such as specific behaviors and complex phenomena like decision-making or emotion [10]. To move beyond the proof-of-concept stage, direct evidence of sonogenetics’ ability to coordinate the modulation of specific neuronal circuits and behavioral outcomes is required. In particular, the circuit-specificity of sonogenetics requires better characterization across different brain regions, given the possibility of confounding elements such as endogenous mechanosensitive ion channels or a non-specific auditory effect triggered by ultrasound. In addition, the temporal resolution of *in vivo* sonogenetics studies needs direct confirmation and validation from selected neurons in response to ultrasound. While the latency of MscL-sonogenetics has been indirectly shown of 150 ms from muscular movement by EMG studies [1], direct profiling of neuronal activity would be important to understand and improve treatment protocols.

Here, we report *in vivo* studies of MscL-mediated sonogenetics to activate specific neurons in intact mouse brains, modulating well-defined neural circuits, and inducing corresponding behaviors in freely-moving mice. We demonstrate the capability of sonogenetics to activate targeted neurons in different subcortical brain regions, including the dorsal striatum (dSTR) and ventral tegmental area (VTA). MscL-sonogenetic stimulation of these regions was found to selectively activate the targeted neural circuits quickly as measured by fiber photometry (FP) and could specifically induce asymmetric locomotion and appetitive conditioning behaviors. Spatial specificity was verified by observing upregulation of the neural activation marker c-Fos and neuronal responses showed sub-second latency. Our functional and behavioral experiments showed that MscL expression significantly enhanced neuronal activation by low-frequency non-invasive ultrasound without the need for intracranial implants or microbubbles.

## Results

### MscL-sonogenetic stimulation of dSTR neurons induces motor response in mice

In a previous study, we used the MscL channel to mediate ultrasound neurostimulation of targeted neurons in the mouse primary motor cortex and dorsomedial striatum [1]. Short ultrasound pulses of 300 ms at low intensities were sufficient to trigger neuronal activation, judged by c-Fos expression and forelimb muscle contractions. Here, we decided to build upon this data and evaluate whether our previously-used protocol was effective and specific enough to also trigger immediate and direct neural activation and corresponding movement behaviors in awake mice. We chose the dSTR as our target because it is a relatively deep region (located ~2.75 mm below the skull) with a well-defined function and behavior pattern, which is relative to initiate and control movement of the body [11–15]. Specifically, it is expected to induce asymmetric locomotion when the neurons are activated.

We examined the direct effect of MscL-sonogenetics on dSTR neuronal activation by measuring fluorescence changes of jRGECO1a (genetically-encoded calcium sensor with red fluorescence) [16] in real-time using FP. Neurons in the right dSTR were simulataneously transduced by hSyn-promoted jRGECO1a and AAVs for EYFP or MscL-EYFP (Fig. 1A). We observed robust expression of EYFP and jRGECO1a colocalized in dSTR neurons (Fig. 1B). Four to five weeks post-transduction, the mice were transcranially stimulated with short US pulses (parameters shown in Fig. 1C) and an optical fiber was used to monitor fluorescence intensity. Prior to US stimulation, both groups showed comparable levels of baseline fluorescence (Peak ΔF/F0 for EYFP = 0.073 ± 0.006%, and MscL = 0.062 ± 0.007%, Fig. 1D, E). However, upon delivery of one 0.15 MPa intensity US pulse, MscL-expressing neurons showed a rapid and significant increase in jRGECO1a fluorescence intensity while EYFP-only neurons did not (Peak ΔF/F0 for EYFP = 0.058 ± 0.008%, MscL = 0.134 ± 0.020%, Fig. 1D). Fluorescence responses to US in the MscL dSTR regions were 2.31-fold of the EYFP dSTR. Acoustic pressures from 0.05 MPa – 0.2 MPa reliably evoked dSTR neural activity in MscL mice, and a generalized pattern of dose-dependence was observed (Fig. 1E). In contrast, the EYFP mice showed a small fluorescence change comparable to spontaneous activity after US stimulation. The latency range of neuronal responses in MscL-expressing neurons was 250.6 ±24.1 ms to 297.3± 31.5 ms (Fig. 1F).

**Fig. 1.**
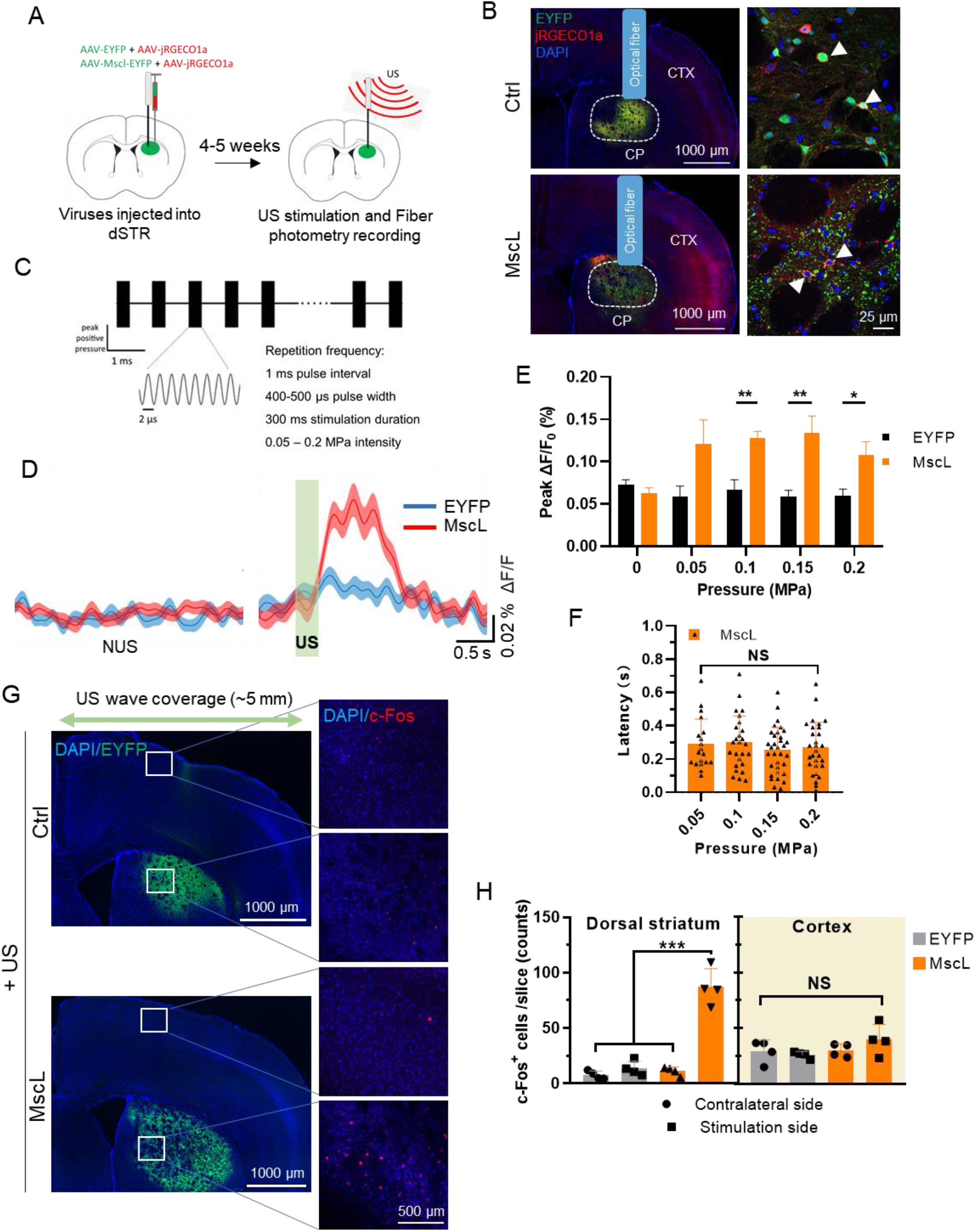
MscL-sonogenetics enables specific neuronal activity in the dorsal striatum. (A) Schematic illustration of *in vivo* real-time calcium activity recording. The right dSTR of mice was co-transduced by AAVs for a calcium sensor (hSyn:jRGECO1a) and either hSyn:MscL-EYFP or EYFP, at a 1:1 ratio. 4-5 weeks later, anesthetized mice were stimulated with US, and neuronal calcium responses were recorded by an optical fiber implanted in the dorsal striatum region. (B) Representative images of dSTR expressing hSyn-EYFP or hSyn:MscL-EYFP and hSyn:jRGECO1a. Caudoputamen (CP), cerebral cortex (CTX). White arrows indicated the EYFP/MscL expressing neurons overlapping with jRGECO1a fluorescence. (C) An illustration of the ultrasound temporal profile used in locomotion experiments. The parameters of ultrasound used were: 0.5 MHz central frequency, 400-500 μs pulse width, 300 ms stimulation duration, 1 kHz PRF, 0.05 – 0.2 MPa intensity ranges. (D) Averaged jRGECO1a fluorescence traces in the dSTR of anesthetized EYFP- or MscL-mice prior to US stimulation (Left) and in response to one 0.15 MPa pressure ultrasound pulse (Right) (0.5 MHz center frequency, 500 μs pulse width, 300 ms stimulation duration, 1 kHz PRF, interval 3s). n=5 mice, 8 trials/each in EYFP group, n=5 mice, 7 trials/each in MscL group. Green rectangle shows the timing of ultrasound stimulation. (E) Average peak Ca^2+^ activity in EYFP- and MscL-mice in response to US pulses of varying intensities (0.05 – 0.2 MPa peak pressure, 500 μs pulse width, 0.5 MHz center frequency, 300 ms stimulation duration, 1 kHz PRF, interval 3s). n = 5 mice in EYFP group, n = 5 in MscL group. * P < 0.05, ** P < 0.01, unpaired t-tests with Holm-Sidak correction. Data are shown as mean ± SEM. (F) Latency between US stimulation (0.05 – 0.2 MPa peak pressure) and detection of an above-threshold fluorescence change. n = 5 in MscL group. NS – not significant, one-way ANOVA with post-hoc Tukey test. Data are shown as mean ± SEM. (G) Representative images of dSTR following US stimulation (0.15 MPa). Images show DAPI + EYFP or MscL-EYFP at low magnification (Left) and DAPI + c-Fos expression in the indicated areas at high magnification (Right). (H) Counts of nuclear c-Fos per slice imaged in the dSTR (Left) and the cortex above targeted region (Right) of mice stimulated with ultrasound. n = 4 mice each group. Data are shown mean ± SEM of average c-Fos+ cells per stained slice. *** P < 0.001, one-way, NS = not significant, ANOVA with post-hoc Tukey test.

To further evaluate the spatial specificity of our scheme, anesthetized mice were treated with US for 40 minutes and their brains were stained for the neural activation marker, c-Fos. Robust EYFP expression was seen in the dSTR of both EYFP and MscL mice (Fig. 1G). Average c-Fos expression in dSTR was found to be significantly higher in the stimulated side of the MscL mice (87 ± 8 c-Fos^+^ cells) than in the contralateral side of the same mice (11 ±2 c-Fos^+^ cells) or in mice expressing only EYFP (stimulation side = 14 ± 3, contralateral = 8 ± 2 c-Fos^+^ cells) (Fig. 1H). Crucially, we found that the cortical regions located directly above the dorsal striatum on the stimulation side did not show a significantly higher level of c-Fos (Fig. 1H). This indicates that our setup could focus the effects of US in the region expressing MscL and not surrounding areas, consistent with our previous results [1]. This is especially important because a single element plane traducer with a diameter of 5 mm was used, meaning that the cortical neurons above striatum were unavoidably sonicated as the ultrasonic waves propagated through. The upregulation of c-Fos being restricted to the region of MscL expression clearly demonstrates the gain-of-function role of the mechanosensitive mediator in sonogenetics. We also stained for c-Fos in the auditory cortex of the brain slices. We did not observe noticeable increase of c-Fos in the auditory cortex in these animals, and found the number of c-Fos in the auditory cortex were not obviously different between EYFP and MscL mice (Supplementary Fig. 1), suggesting that auditory pathway did not play a major role in long term effects. Thus, MscL-sonogenetics could specifically activate neurons in the dSTR.

We next tested whether the MscL-sonogenetics was effective enough to modulate the expected asymmetric locomotion in freely-moving mice. A wearable US transducer was attached above the virally-transduced dSTR in freely moving mice and their locomotion was recorded during US stimulation (Fig. 2A, B). Robust EYFP or MscL-EYFP signals were observed in the right dSTR region (Fig. 2C) confirming successful viral expression. Mice were then treated with ultrasound in an open field box and their movement behaviors were recorded and analyzed. Ultrasound stimulation consisted of three 1 minute epochs (pre-stimulation (“Pre”), ultrasound stimulation (“US”), post-stimulation (“Post”). MscL-expressing mice were seen to obviously increase their locomotor activity during ultrasound treatment, and returning to their baselines post-US, compared to negligible change for EYFP mice (Fig. 2D). The mobility speed of the MscL mice increased from 66.4 ± 3.4 mm/s to 83.8 ± 2.3 mm/s, while the EYFP mice did not show obvious change (Fig. 2E, F). MscL moved greater distances during the periods of US stimulation than EYFP mice (distance during US: MscL = 3776.0 ± 263.3 mm, EYFP = 1753 .0± 179.6 mm), and all mice reduced their activity in the periods between rounds of stimulation (Fig. 2G,H). EYFP mice also showed small increases in their motor activity during US stimulation, but the magnitude of these changes was small and not significant compared to the pre-US measurements. Thus, MscL-sonogenetics in the dSTR could effectively induce significant motor responses of awake mice, consistent with previous optogenetic neuromodulation targeting the same region [13, 17].

**Fig. 2.**
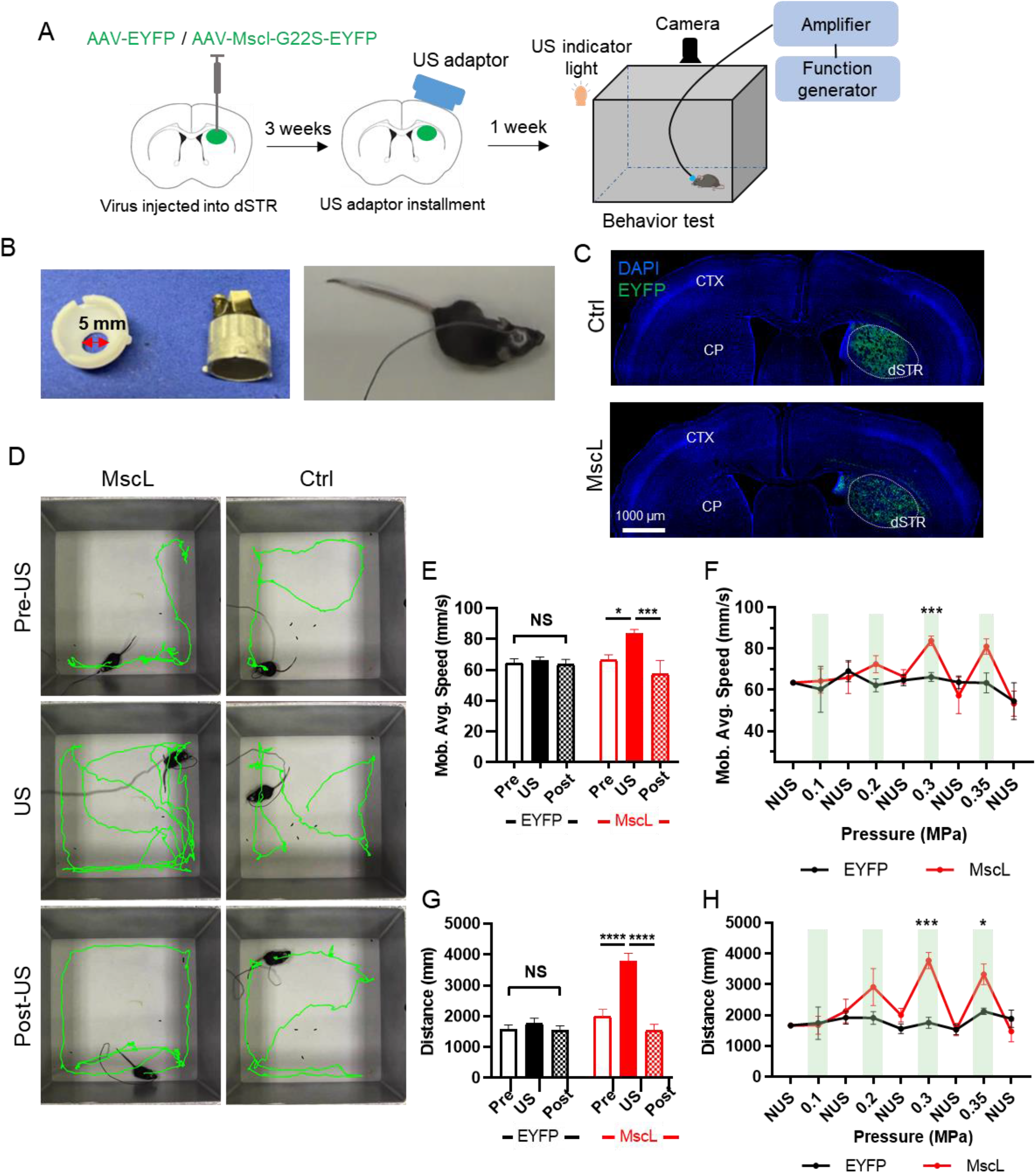
Ultrasound stimulation evokes significant increase in locomotion behavior in mice with MscL-expressing dorsal striata. (A) Schematic illustration of locomotion experiments. Briefly, mice were injected with hSyn:EYFP or hSyn:MscL-EYFP in their right dorsal striatum. Three weeks later, an ultrasound adaptor was installed. One week later, mice were placed in an open field box with their movement recorded before, during and after ultrasound stimulation. The behavior documented in the video was then analyzed and quantified. The parameters of ultrasound used were: 0.9 MHz central frequency, 400 μs pulse width, 300 ms stimulation duration, 1 kHz PRF, 0.1 – 0.35 MPa intensity ranges. (B) Images of the ultrasound adaptor and the wearable ultrasound transducer (Left). The diameter of the ultrasound transducer was 5 mm. The wearable ultrasound transducer was mounted on the adaptor attached onto the mouse head (Right). (C) Representative images showing expression of DAPI + hSyn:EYFP (Top) or hSyn:MscL-EYFP (Bottom) in the dorsal striatum (dSTR). Caudoputamen (CP), cerebral cortex (CTX). (D) Representative trajectories recorded from mice stimulated in the dorsal striatum pre-US, during US, and post-US with 0.3 MPa ultrasound application (each trace 1 min long). (E) Comparison of mobility average speed of the EYFP and MscL-expression mice, before, during and after US stimulation (0.3 MPa). n = 5 mice in EYFP group, n = 6 mice in MscL group. Data are shown as mean ± SEM; *p < 0.05, *** P < 0.001, NS = not significant, two-way ANOVA with post-hoc Tukey test. (F) Summary of mobility average speed of EYFP and MscL-expression mice with stimulation of different ultrasonic intensities from 0.1 – 0.35 MPa peak pressure. Green bars indicate the timing of ultrasonic stimuli. Data are shown as mean ± SEM; * P < 0.05 and *** P < 0.001, unpaired t-test with Holm-Sidak correction. (G) Comparison of the horizontal distance covered by the same mice as in (E). n = 5 mice in EYFP group, n = 6 mice in MscL group. Data are shown as mean ± SEM; **** P < 0.0001, NS = not significant, two-way ANOVA with post-hoc Tukey test. (H) Summary of distance of EYFP and MscL-expression mice with stimulation of different ultrasonic intensity from 0.1 – 0.35 MPa peak pressure. Green bars indicate the timing of ultrasonic stimuli. Data are shown as mean ± SEM; * P < 0.05 and *** P < 0.001, unpaired t-test with Holm-Sidak correction.

Taken together, our findings suggest that MscL-sonogenetics can specifically stimulate neurons in the dSTR and induce characteristic asymmetric locomotion behavior.

### Efficient dopamine release and appetitive conditioning by MscL-sonogenetics modulation of VTA reward circuitry

We next tested the feasibility of using MscL-sonogenetics to control higher-level behavior by targeting more specific neurons. The VTA was chosen as it is a midbrain reward center where the dopaminergic circuit is well-defined and involved in various reward related behavior and diseases [18]. In order to test whether MscL-sonogenetics could target specific circuits, dopaminergic (DA) VTA neurons were selectively made to express MscL. We used a dual-viral vector strategy: one vector delivered Cre recombinase under the modulation of a tyrosine hydroxylase promoter (TH-promoter) [19, 20], the other delivered a Cre-recombinase-dependent EYFP or MscL-EYFP fragment (Fig. 3A). Confocal imaging confirmed that TH^+^ neurons overlapped with EYFP or MscL-EYFP in the VTA region (Fig. 3B) indicating successful expression of MscL in dopaminergic neurons after 4-5 weeks.

**Figure 3.**
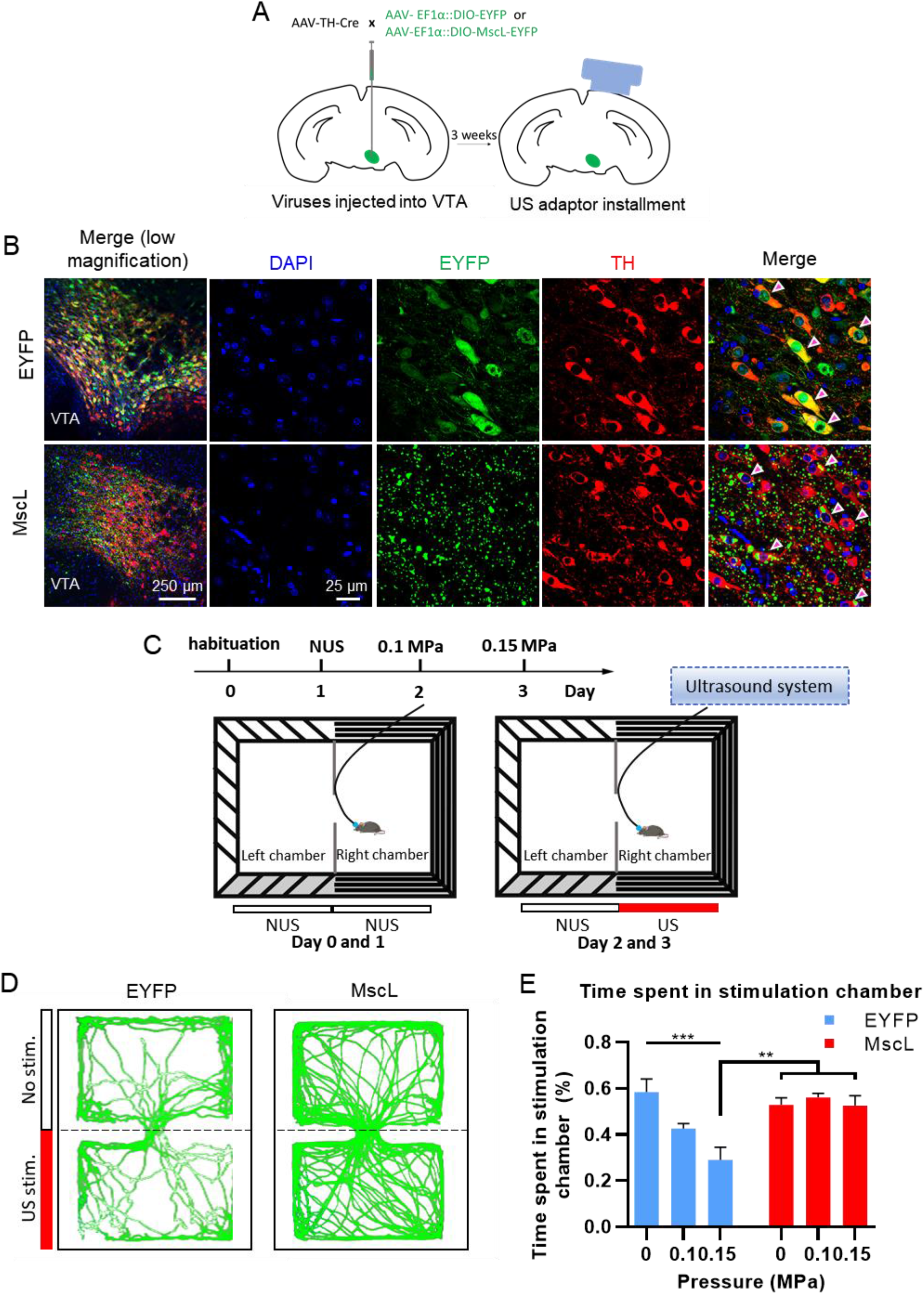
Selective activation of mesolimbic pathway by MscL-sonogenetics enables appetitive conditioning. (A) Schematic illustration of the transduction protocol for dopaminergic neuron-specific transduction of MscL-EYFP or EYFP. Briefly, mice were injected with a mix of AAV-TH-Cre and AAV-EFlα::DIO-EYFP or AAV-EFlα::DIO-MscL-EYFP in the right ventral tegmental area. Three weeks later, an ultrasound adaptor was installed followed by one week for recovery, then the mice were stimulated with ultrasound. (B) Representative images depicting VTA regions expressing TH:EYFP (Top) or TH:MscL-EYFP (Below). Arrows denote example EYFP+/TH+ neurons. (C) Schematic illustration of the real time place preference assay. A two-chambered box was prepared, one chamber having horizontal stripes and the other with vertical stripes. Mice were habituated and tested according to the protocol illustrated. The movement of mice between chambers was recorded by a digital camera, with US stimulation being assigned to a random chamber. US stimulation was timed to start and stop immediately when a mouse entered and exited the stimulation chamber respectively. (NUS = no ultrasound) (D) Representative path-tracing of a mouse with TH-cre-EYFP (control) or TH-cre-MscL-EYFP expression during the real-time place preference test (0.1 MPa). Red bar indicates stimulation side. (E) Calculated percentage of time spent by mice on the stimulation side with NUS, 0.1 MPa or 0.15 MPa US in EYFP and MscL mice. n =5 mice in EYFP mice; n = 5 mice in MscL-expressing mice. ** P > 0.01, ** P > 0.01, two-way ANOVA with post-hoc Tukey test. Data are shown as mean ± SEM.

We tested whether MscL-sonogenetics could specifically modulate appetitive conditioning using the real-time place preference assay. Habituated mice were placed in a rectangular open field box with two distinct chambers, one paired with US stimulation, the other paired with non-US stimulation (NUS), and mice could move freely between chambers (Fig. 3C). Mice were wearing a transducer aimed at the VTA (Fig. 3A). US stimuli were triggered immediately when a mouse entered the US chamber and turned off immediately when the chamber was exited. US stimulation induced aversive responses with 0.1 MPa US in EYFP mice (Fig. 3D, E), consistent with previous reports [21]. Increasing acoustic pressure to 0.15 MPa in EYFP mice further aversive behavior (Fig. 3E). In contrast, we found that MscL mice showed a significant preference for the US chamber compared to EYFP mice. MscL mice treated with 0.1 and 0.15 MPa US spent a significantly higher proportion of time in the in the stimulation chamber than EYFP mice treated with the same US pressure (Fig. 3D,E). Furthermore, we found that DA neuron-specific sonogenetic stimulation in VTA could activate neuronal population in the NAc in acute *ex vivo* brain slices (Supplementary Fig. 2 A,B). NAc neurons displayed calcium fluorescence increases time-locked with MscL-expressing VTA stimulated with US (Supplementary Fig. 2C), and responses from MscL-neurons were significantly higher than EYFP-neurons (Supplementary Fig. 2D). This suggests that MscL-sonogenetics could stimulate dopaminergic circuits in the VTA and induce the reversal of aversive responses. In summary, we found that US stimulation of mouse of MscL-expressing dopaminergic neurons in the VTA could successfully and specifically induce appetitive conditioning behavior, reversing obvious aversive responses seen from mice without MscL. Thus, the MscL-sonogenetics setup could effectively enable control reward behavior.

Finally, to confirm whether MscL sonogenetics could actually activate VTA reward circuitry, we recorded downstream dopamine release in the NAc [22, 23]. Neurons in the ipsilateral NAc were transduced with the fluorescent dopamine sensor DA2m [24] under an hSyn promoter. An optical fiber was inserted into the NAc to monitor dopamine activity through the sensor’s fluorescence change (Fig. 4A). We found robust DA2m expression in NAc region (Supplementary Fig. 3A) and MscL / EYFP signal in DA-neurons of the VTA (Supplementary Fig. 3B). Upon stimulation of the VTA with pulses of 0.3 MPa ultrasound, we found rapid synchronous increases in DA2m fluorescence in the NAc of the MscL mice condition but not EYFP mice (Peak ΔF/F0, EYFP = 0.23%, MscL = 0.52%) (Fig. 4B). NUS and EYFP+US conditions at both tested US intensities showed very little fluorescence change, and the MscL + US conditions were larger in magnitude at both 0.1 MPa (Peak ΔF/F0, EYFP = 0.18%, MscL = 0.41%) and 0.3 MPa (Peak ΔF/F0, EYFP = 0.23%, MscL = 0.52%), although only the 0.3 MPa condition was statistically significant (Fig. 4C). DA signal responses to US stimuli were found to be repeatable and with a low latency of ~ 250 ms at both tested US intensities (Fig. 4D,E). Taken together, this suggests that the avoidance behavior displayed by EYFP mice could be due to non-specific stimulation of the VTA, unrelated to reward circuits, while MscL-sonogenetics could specifically induce DA secretion in neurons projecting to the NAc *in vivo*, eliciting place preference behavior.

**Figure 4.**
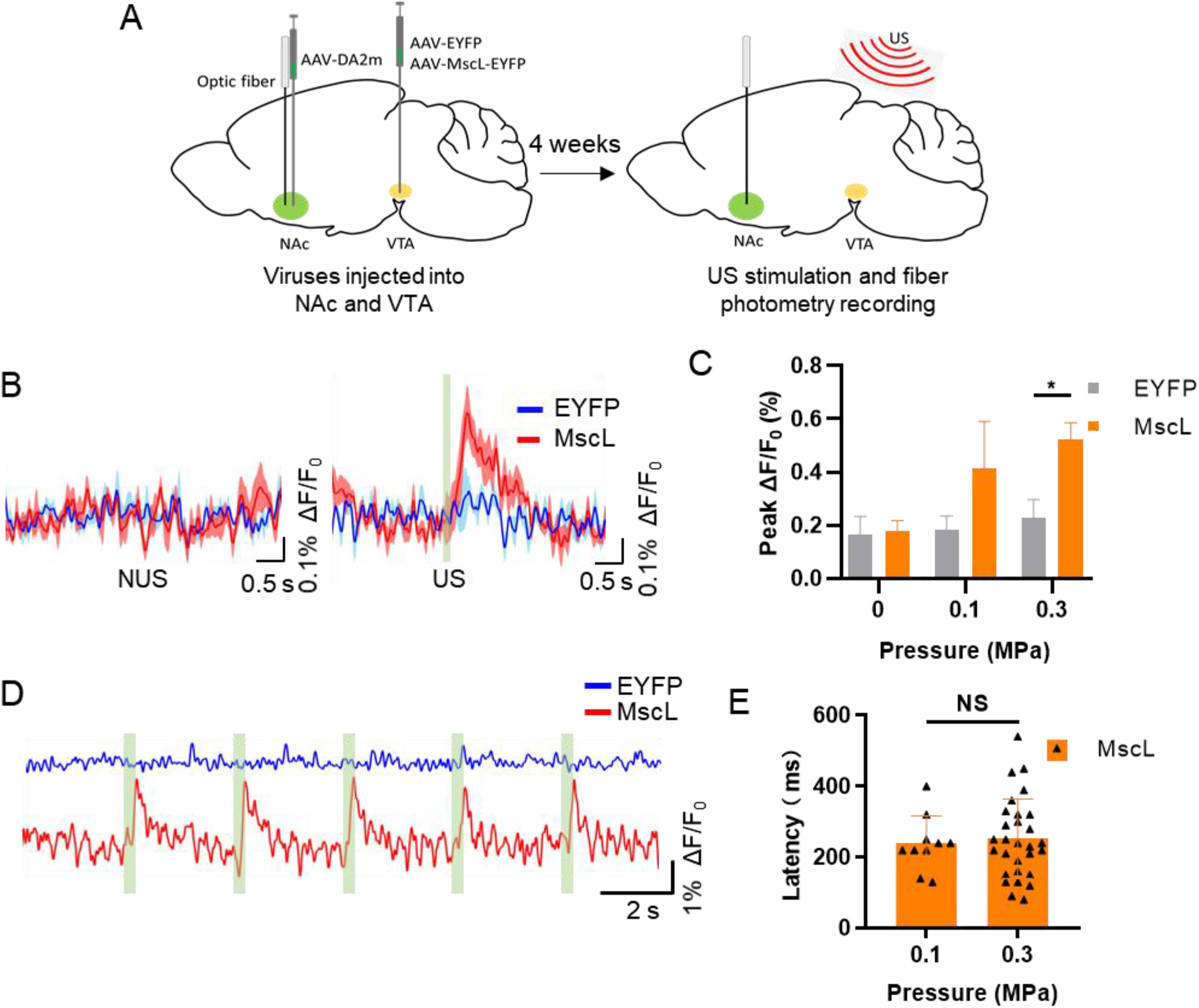
Efficient sonogenetically-enabled dopamine release in the NAc through modulation of the mesolimbic pathway. (A) Schematic illustration of dopamine fiber photometry experiments. Mice were injected with AAV-hSyn-DA2m in right hemisphere of NAc and hSyn-EYFP or hSyn-MscL-EYFP in the ipsilateral VTA. After viral injection, an optical fiber was inserted into NAc. Four weeks later, mice were treated with US and dopamine fluorescence signals were recorded. (B) Averaged DA2m fluorescence signal change (ΔF/F_0_) in the NAc of anesthetized EYFP- and MscL-expressing mice, with or without 0.3 MPa US stimulation. Blue traces show DA signals in the EYFP-mice, while red traces show the DA signals in MscL-expressing mice. Green rectangle shows the timing of ultrasound stimulation. n=4 mice, 6 trials/each in EYFP group. n=5 mice, 6 trials/each in the MscL group. (C) Average peak DA2m activity (ΔF/F_0_) in response to 0, 0.1, 0.3 MPa ultrasound stimulation (0.5 MHz center frequency, 500 μs pulse width, 300 ms stimulation duration, 1 kHz PRF, interval 3s) in EYFP- and MscL-expressing mice. n=4 mice, 6 trials/each in EYFP group. n=5 mice, 6 trials/each in the MscL-expressing group. * P < 0.05, **, unpaired t-tests with Holm-Sidak correction. Data are shown as mean ± SEM. (D) Representative DA2m fluorescence traces in the NAc of anaesthetized EYFP and MscL-expressing mice when stimulated by multiple pulses of 0.3 MPa US. (E) Latency between US stimulation and detection of an above-threshold fluorescence change in MscL mice. Data are shown as mean ± SEM. NS = not significant, unpaired two-tailed t-test.

## Discussion

The present study demonstrates that MscL-mediated sonogenetics can stimulate specific neural circuits effectively with high spatiotemporal precision, providing a non-invasive way to modulate higher behaviors in mouse models. We confirmed that MscL-sonogenetics can target a neural circuit to promote specific signaling and behaviors. In addition, we show that MscL-sonogenetics could induce rapid responses to US stimuli through FP recordings in deep brain regions. The latency of fluorescence change was consistent between the genetically-encoded calcium and dopamine sensors in the two regions, at ~250 ms. Although the temporal resolution of sonogenetics could not be determined by patch clamp, due to its incompatibility with ultrasonic vibrations, the quick responses measured by fiber photometry suggest a sub-second temporal precision. Finally, MscL-sonogenetics could elicit well-defined behaviors through specific activation of established neural circuitry. Our stimulation scheme showed good spatiotemporal resolution when targeted to brain regions at different depths, without obvious non-specific activation and responses closely synchronized with US stimuli.

Sonogenetics has the advantage of combining a modular element (the choice of mechanosensitive mediator) and a non-invasive stimulation modality (ultrasound). Here, we were able to further increase the specificity of the treatment by injecting virus in a specific brain region, and using the Cre-Lox system to preferentially express MscL only in certain cell types. This combinatory approach offers multiple opportunities to tailor the treatment for the intended application, and the prospect of continuous improvement. Choosing a different transducer or ultrasound parameters, different brain region or a different cell-type for targeting could allow the study of other aspects of brain function. The discovery and development of other mediating molecules, such as other mechanosensitive ion channels [6] or non-channel transmembrane proteins [3], could provide new ways to stimulate desired regions with ultrasound. It could also be possible to further reduce the minimal invasiveness of the treatment by making use of new genetic technologies. For instance, the AAV-PHP.eB virus has been shown to be capable of transducing cells in the brain through mere tail-vein administration [25, 26]. Such a system could be used to preferentially transduce targeted cells in the brain while also possibly reducing the need to deliver the virus directly into the brain. Finally, we found MscL-sonogenetics to be compatible with fiber photometry and free-moving recording without significant interference and crosstalk. Thus, integration of these important techniques could help for comprehensive study of the brain and how it works. For instance, sonogenetics could further be used with fMRI, a well-established technique, and could thus enable screening and dissection of neural circuits in large animals, e.g. non-human primates. Thus, sonogenetics offers an adaptable, modular approach that is amenable to integration with other modalities, which could allow for wider-ranging study.

In addition to the modularity and non-invasiveness of the approach, another advantage of sonogenetics is that it can affect mesoscale circuits in the brain. Optogenetics has been used to dissect brain microcircuits affecting a specific brain function [27]. However, understanding how these microcircuits interact, and how higher-order functions are constituted would require modulation of mesoscale circuits. As shown here, with a flat single-element ultrasound transducer, MscL-sonogenetics can induce cell-type-selective stimulation in the midbrain and deep brain dependent on the spatial profiles of MscL expression. Given the dynamic focusing capability, sonogenetics could provide an opportunity to stimulate mesoscale circuits non-invasively with defined spatiotemporal pattern to unlock the complex functions and dysfunctions.

A potential concern about sonogenetics is the activation of neurons not expressing the chosen mediator, causing background responses and compromising specificity and efficacy. Ultrasound has been shown to activate various endogenous mechanosensitive ion channels, including Piezo1 [28, 29], TRAAK [4], TRPA1 [6] etc. We tried to minimize non-specific effects by using short US pulses at the lowest effective acoustic pressure. Our application of sonogenetics in deeper brain regions, e.g. VTA and dSTR, did not show obvious off-target effects in the present study, suggesting that MscL-sonogenetics is capable of stimulating specific neural circuits. However, more detailed characterization of sonogenetics’ performance in other brain regions remains critical, especially in the cortex, as an auditory confound is a perennial concern for US treatment [30].

We envision that an approach like sonogenetics could be promising for the eventual development of neuropsychiatric treatments. For instance, we found that MscL-sonogenetics could be used to induce dopamine secretion through the mesolimbic pathway. Impaired dopamine transmission in the human brain is related to neurological and psychiatric diseases, such as Parkinson’s disease [31, 32], schizophrenia [33] and depression [34]. Unlike pharmacological or lesion-based interventions, sonogenetics is non-invasive with cellular specificity and neural circuit accuracy. Regulating dopamine dynamics in the targeted region, might help alleviate symptoms of such neurological diseases. In addition to dopamine, this approach may also regulate other neurotransmitters by manipulating specific neurons in appropriate brain regions, such as glutamatergic or GABAergic neurons. Thus, we believe that sonogenetics approach has the potential to become a novel alternative approach for treating neurological and psychiatric diseases.

## Acknowledgements

This work was supported by the Hong Kong Research Grants Council General Research Fund (15104520, 15102417 and 15326416), Hong Kong Innovation Technology Fund (MRP/018/18X and MHP/014/19), Key-Area Research and Development Program of Guangdong Province (2018B030331001), and internal funding from the Hong Kong Polytechnic University (1-ZE1K, 1-BBAU and 1-ZVW8). The authors would like to thank the facility and technical support from University Research Facility in Life Sciences (ULS) and University Research Facility in Behavioral and Systems Neuroscience (UBSN) of The Hong Kong Polytechnic University.

## Author contributions

Conceptualization, Q.X., Z.Q. and L.S.; Methodology, Q.X., Z.Q., J.G. and L.S.; Investigation, Q.X., S.M., K.F.W. J.Z., X.H., W.Y. and S.K.; Data Analysis, Q.X., S.M., X.H., Z.Q., J.G. and S.K.; Manuscript Preparation, Q.X., Z.Q., S.K. and L.S.; Funding Acquisition, L.S.

## Materials and Methods

### Animal subjects

Male, 6-8 weeks old, C57BL/6J mice, were used for this study. Mice were housed under standard conditions with food and water available ad libitum. Mice were habituated to the procedure room for at least 30 min prior to behavioral testing experiments. All animal experiments were approved by the Animal Subjects Ethics Sub-Committee (ASESC) of the Hong Kong Polytechnic University. Animal use and care were performed following the guidelines of the Department of Health - Animals (Control of Experiments) of the Hong Kong S.A.R. government.

### Stereotaxic injection

Mice were anesthetized with ketamine and xylazine (100 mg/kg and 10 mg/kg respectively), and placed in a stereotaxic instrument (RWD, Shenzhen, China). A small craniotomy hole was made over the targeted area. Viral vectors were micro-injected into the right side of mice brain by standard stereotaxic procedures [35]. AAVs were injected at the rate of 0.05 – 0.1 μl per minute. The micro-syringe was left in place for an extra 10 min before withdrawal.

For locomotion experiments, 500 nl of AAV-hSyn-EYFP or AAV-hSyn-MscL-G22S-EYFP (BrainVTA (Wuhan) Co. Ltd) was unilaterally injected into the dorsal striatum at AP +0.50 mm, ML −1.8 mm, from Bregma; and DV −2.75 mm from the brain surface. Three weeks post-transduction, a portion of the scalp was excised at the above coordinates, and an ultrasound adaptor was fixed to the skull using dental cement.

For calcium fiber photometry in the dorsal striatum, mice were injected with virus vectors in the right side of dorsal striatum (DV: −2.75mm) as mentioned above. The volume ratio of the viral vector mixture was 1:1 for AAV-hSyn-jRGECO1a (OBiO Technology, Shanghai) and AAV-hSyn-EYFP or AAV-hSyn-Mscl G22S-EYFP in dorsal striatum. After micro-syringe withdrawal, an optic fiber was inserted into the location of the syringe.

For dopamine fiber photometry in the VTA, 300 nl AAV-hSyn-EYFP or AAV-hSyn-Mscl-G22S-EYFP were micro-injected into the right side VTA (AP: - 2.9 mm, ML: −0.5 mm, DV: −4.5 mm), and 1 μl hSyn-DA2m (Vigene Bioscience Co., Ltd) were injected into the ipsilateral side of nucleus accumbens (NAc, coordination: AP: 1 mm, ML: −1.1 mm, DV: −4.1 mm). An optical fiber was implanted in the NAc region after AAV injection.

For reward-related experiments (real time place preference), 300 nl AAV-EF1a::DIO-EYFP or AAV-EF1a::DIO-MscL-G22s-EYFP (BrainVTA (Wuhan) Co. Ltd) mixed with pAAV-TH-Cre-WPRE-hGHpA (OBiO Technology, Shanghai) (1:1) were micro-injected into the right side VTA (AP: - 2.9 mm, ML: −0.5 mm, DV: −4.5 mm). An ultrasound adaptor was installed above the targeted region after 3 weeks of viral transduction.

For the VTA and NAc acute brain slice calcium imaging experiment, 300 nl AAV-EF1a::DIO-EYFP or AAV-EF1a::DIO-MscL-G22s-EYFP mixed with pAAV-TH-Cre-WPRE-hGHpA (1:1) were micro-injected into the right side VTA (AP: - 2.9 mm, ML: −0.5 mm, DV: −4.5 mm) and 0.5 μl hSyn-jRGECO1a were injected into the ipsilateral side of NAc (AP: 1 mm, ML: −1.1 mm, DV: −4.1 mm).

### Fiber photometry recording

Fiber photometry recordings were performed at least 4 weeks after viral injection. Mice (jRGECO1a- or DA2m-expressing mice) were anesthetized with 1-2% isoflurane and the hair was removed with scissors. Eye ointment was applied to prevent corneal drying. The implanted fiber was connected to fiber optic meter (Thinker Tech Nanjing BioScience Inc) through an optical fiber patch cord for guiding the light. Fiber photometry recoding was performed using a 570 nm laser at around 50 μW for jRGECO1a, and a 480 nm laser at around 50 μW for DA2m. Data was collected at 100 Hz and analyzed using a customized MATLAB script. The fluorescence change (ΔF/F0) was calculated as (F-F0)/F0, where F0 is the baseline fluorescence signal.

### Open field recording of locomotion

1 week after adaptor installation, a 0.9 MHz wearable ultrasound transducer was plugged into it. Mice were placed into the center of an open field chamber (40 cm length × 40 cm width × 30 cm height). Mice were habituated in the chamber for 15-20 min prior to ultrasound stimulation. Mouse movements were then captured by a digital camera (Canon, LEGRIA, HF, M506) placed at ~ 45 cm above the open field chamber. Mice were allowed to rest 1-2 min between each trial. The total travel distance and the mobility speed of mice were calculated from the recorded video clips using ToxTrac software [36, 37] and normalized to that of the same distance / mobility speed on the first average measurement. A mouse was considered to be moving at a minimum speed of 20 mm/s.

### Real time place preference assay

A 50 × 60 cm arena was divided into left and right chambers of equal size (50 cm length × 30 cm width × 30 cm height). The walls on each side were marked with horizontal stripes or vertical stripes. Mice were allowed to habituate in the arena on day 0. Mice were then stimulated with the ultrasound apparatus at 0, 0.1 or 0.3 MPa per daily session until completion of the experiment. Mice were allowed to freely explore the two chamber arenas for 10 min. One chamber was randomly assigned as being the initial stimulation chamber. When a mouse entered the stimulation chamber, ultrasound delivery was immediately begun, until it crossed over to the non-stimulation chamber, at which point the US was immediately stopped. The movement of the mice was recorded via a digital camera and the time of the mice spent in each chamber (stimulated and non-stimulated) was calculated by ToxTrac software. Data from mice with a strong initial preference or avoidance for either chamber (by spending >75% or <25% of test time in either chamber) were discarded to ensure an unbiased outcome [38].

### Immunohistochemical fluorescent staining of c-Fos

90 minutes after ultrasound treatment, mice were perfused with 0.9% saline, followed by 4% paraformaldehyde (PFA) (cat. no. P1110, Solarbio). After dissection, brains were post-fixed overnight in 4% PFA. Coronal sections were prepared from the brain at a thickness of 40 μm by a vibratome. Slices were blocked using blocking buffer (10% normal goat serum + 1% BSA + 0.3% Triton in 1X PBS) and incubated overnight in primary antibody solution diluted in the blocking buffer. Slices were then washed with PBS 3 times, incubated with secondary antibodies diluted in blocking buffer for two hours at room temperature. Slices were then washed 3 times, coverslips dried, and mounted on glass slides using small drops of Prolong Diamond Antifade Mountant with DAPI. Primary antibodies used were c-Fos (2250, Cell Signaling Technology, dilution 1:500), Tyrosine hydroxylase (MAB318, Millipore, dilution 1:500). Secondary antibodies used were goat anti-rabbit IgG (H+L), Alexa Fluor 555 (A-214428, Invitrogen, dilution1:1,000) or goat anti-mouse IgG (H+L), Alexa Fluor 555 (A-32727, Invitrogen, dilution 1:1000). Each sample was divided into 3 sets and 1 set was used to stain for c-Fos expression. Each set contained 5-8 brain slices. The number of cells showing c-Fos, were counted using ImageJ, and the number of c-Fos^+^ cells per 733 x 733 μm slice were calculated. All brain slices were imaged using the confocal microscope (TCS SP8, Leica) in the ULS facilities in The Hong Kong Polytechnic University.

### Preparation and fluorescence imaging of acute brain slices

4 weeks after virus injection, mice were anesthetized with an intraperitoneal injection of Ketamine and Xylazine (100 mg/kg and 10 mg/kg respectively) and perfused with ice-cold oxygenated NMDG ACSF buffer containing (in mM): 92 N-methyl-D-glucamine (NMDG), 2.5 KCl, 1.25 NaH2PO4, 20 NaHCO3, 10 HEPES, 25 Glucose, 2 thiourea, 5 Na-ascorbate, 3 Na-pyruvate, 0.5 CaCl2, 10 MgSO2, 12 NAc. The pH of the ASCF was 7.3-7.4. The brains were immediately removed and placed in ice-cold oxygenated slicing buffer. The brains were sectioned into 300 um thick slices using vibratome (Leica VT1200), and the slices were incubated at 34 °C for at NMDG ACSF for 10-12 min, followed by N-2-hydroxyethylpiperazine-N-2-ethanesulfonic acid (HEPES) ACSF that contained (in mM): 92 NaCl, 2.5 KCl, 1.25 NaH2PO4, 30 NaHCO3, 20 HEPES, 25 Glucose, 2 thiourea, 5 Na-ascorbate, 3 Na-pyruvate, 2 CaCl2, 2 MgSO2 for at least 1 h at 25 °C. The brain slices were transferred to a slice chamber for calcium fluorescence imaging and were continuously perfused with standard ACSF that contained (in mM): 119 NaCl, 2.5 KCl, 1.25 NaH2PO4, 24 NaHCO13, 1.25 Glucose, 2 CaCl2, 2 MgSO2 at 2-3 ml/min. An ultrasound transducer with a plastic tube was placed above the VTA region, the neural activity of NAc neurons were recorded by a modified inverted epifluorescence microscope. The fluorescence was recorded using a 575-645 nm filter.

### Ultrasound instrument and transducer installation

To set up the electronic system driving the ultrasound transducer, the output of a function generator (AFG251, Tektronix) was first connected to the input of a power amplifier (A075, Electronics & Innovation Ltd) with a BNC wiring, then the output of the amplifier was connected to an ultrasound transducer (I7-0012-P-SU, Olympus or wearable transducer). The skull above targeted region was cleaned with saline cotton. The wearable ultrasound transducer was mounted upon the adaptor attached to the skull, coupled with ultrasound gel.

### Data processing

The counting of c-Fos^+^ cells was single-blinded, performed by an experimenter who did not know the groups beforehand.

Behavioral tests data analysis was done by experimenters blinded to experimental conditions using ToxTrac software.

### Statistical analysis

All statistical analyses were performed using the GraphPad Prism software. Two-tailed unpaired t-tests, one- or two-way ANOVA, with post-hoc Tukey or Holm-Sidak test as appropriate, were performed to determine statistical significance. Summary results were presented as mean ± SEM. P values below 0.05 were considered statistically significant.

